# An osmolarity dependent mechanism partially ameliorates retinal cysts and rescues cone function in a mouse model of X-linked Retinoschisis

**DOI:** 10.1101/2023.10.09.561513

**Authors:** Ella J. Gehrke, Jacob Thompson, Emily Kalmanek, Sarah Stanley, Sajag Bhattarai, Brianna Lobeck, Sara Mayer, Angela Mahoney, Salma Hassan, Ying Hsu, Arlene V. Drack

## Abstract

**Introduction:** X-linked retinoschisis (XLRS) is a vitreoretinal dystrophy caused by *RS1* gene mutations which disrupt retinoschisin protein function. A vital protein for maintaining retinal architecture, the absence of functional retinoschisin leads to the development of intraretinal cysts. The preliminary goal of this study was to investigate a low dose gene therapy in *Rs1* knockout (*Rs1*-KO) mice; however, our experiments revealed an unexpected therapeutic effect of a hypertonic buffer, which led to further exploration of this effect.

**Methods:** 10 *Rs1-*KO mice were subretinally injected with an AAV2/4 vector containing the *RS1* gene driven by an *Ef1α* promoter. 16 *Rs1-*KO mice were subretinally injected with a hypertonic buffer (180 mM NaCl 0.001% F68/PBS (pH 7.4)) or an isotonic buffer (155.2 mM NaCl 0.001% F68/PBS, pH 7.0) as a sham control. Endpoints included electroretinogram (ERG), optical coherence tomography (OCT), and a visually guided swim assay (VGSA). An immunohistochemistry assay was used to quantify cone density in buffer injected and treatment-naïve eyes.

**Results:** Unexpectedly, hypertonic buffer-injected eyes had significantly reduced cyst severity at 1 month post-injection (MPI) (*p=<0*.*0001*), significantly higher amplitudes in cone-dominant ERGs persisting to 5 months post-injection (*5 Hz flicker; p=0*.*0018; 3*.*0 Flash; p=0*.*0060*) and demonstrated improved navigational vision in the light compared to untreated *Rs1*-KO eyes (*p<0*.*0001*). To investigate the role of tonicity on this effect, an isotonic buffer-injected cohort was created (155.2 mM NaCl 0.001% F68/PBS, pH 7.0) (n=6). Surprisingly, hypertonic buffer-injected eyes exhibited a greater reduction in cyst severity and demonstrated improved cone-dominant ERG metrics over isotonic buffer-injected eyes. Using an immunohistochemistry assay, we demonstrated greater cone density in hypertonic buffer-injected eyes than untreated controls (*p=0*.*0147)*, suggesting a possible cone preservation mechanism. Moreover, our findings reveal a negative correlation between the peak severity of cysts and long-term cone-dominant ERG metrics, implying that effectively managing cysts could yield enduring benefits for cone function.

**Discussion/Conclusion:** This study presents evidence that cyst resolution can be triggered through an osmosis-dependent pathway, and cyst resolution can have long term effects on cone signaling and survival, offering potential insights for the development of novel treatments for patients with XLRS.

## 1 Introduction

Juvenile X-linked Retinoschisis (XLRS), a vitreoretinal disorder, is the leading cause of macular dystrophy in young males. Its prevalence on a global scale reaches up to 1 in 5,000 individuals (1). The pathology of the disease involves an X-linked autosomal recessive mutation in the retinoschisin 1 (*RS1)* gene located at Xp22.1-p22.3 (1). Mutations in *RS1*, of which there are greater than 200 known, impair production of the 224 amino acid protein, retinoschisin (1).

Retinoschisin is primarily secreted by photoreceptors and bipolar cells, and influences cell-cell interactions through a ubiquitous discoidin domain, facilitating photoreceptor-bipolar cell signaling and stabilizing the extracellular environment of the retina (2, 3). In the absence of a functional retinoschisin protein, the laminar architecture of the retina becomes disrupted, causing retinal layer separation, known as schisis, and the formation of cysts. Cysts, a hallmark of XLRS, spatially divide the inner nuclear and outer plexiform layers, separating the bipolar cells and photoreceptors.

From a clinical perspective, essentially all patients with XLRS experience foveal involvement, impairing central vision (4). Comparatively, around 50 percent of patients also exhibit peripheral involvement (2, 4). The progressive impairment of central vision due to macular atrophy can result in substantial functional vision loss. Additionally, the development of schisis significantly increases the risk of vitreous hemorrhage and full thickness retinal detachment following even mild ocular trauma (4). Strategies to manage cyst severity are needed.

Currently, Carbonic Anhydrase II Inhibitors (CAIs) serve as the primary medical therapy available for patients with XLRS. The proposed mechanism of CAIs involves inhibition of membrane-bound carbonic anhydrase (CA) in the retinal pigment epithelium (5). Fluid transport from the subretinal space into the choroid is enhanced experimentally following introduction of a carbonic anhydrase inhibitor (CAI), acetazolamide, suggesting CA is involved in maintaining fluid homeostasis of the retina (5). However, despite their efficacy in reducing intraretinal cyst formation in some patients, CAIs are associated with well-known systemic side effects affecting the endocrine, gastrointestinal, cardiovascular, and central nervous system, potentially impacting medication compliance (6-8).

While CAIs have shown short term benefits including mild decrease in central macular thickness, reduction of intraretinal cyst formation, and improved vision, it is not known whether they prevent the long-term progression of XLRS (7). Sudden discontinuation of brinzolamide, a commonly used CAI in clinical practice, can lead to increased cyst formation and elevated intraocular pressure during periods of drug holiday (7, 8). There are currently no studies demonstrating long-term efficacy or safety. In addition, it is not clear why reducing cyst severity alone, without rescuing the *RS1* gene, is associated with vision improvement.

Given the lack of effective long-term therapy, ongoing efforts in gene therapy have emerged as a potential avenue for treatment. A Phase I/IIa trial was conducted on human subjects which involved a single intravitreal injection of an AAV8 vector carrying the *RS1* gene (9). This safety study was introduced after good success in a mouse model; however, in human subjects, several ocular inflammatory outcomes were recorded (9). Additional human studies have also noted inflammatory outcomes in participants receiving intravitreal high dose AAV gene therapy for XLRS (10).

Compared to intravitreal injections delivering equivalent doses of AAV, subretinal injections elicited lower levels of humoral immune reactions both in mice and in primates (11). In our study, we investigate whether a subretinal injection of low-dose adeno-associated virus, AAV2/4-EF1α-*RS1*, can rescue the retinal phenotype with reduced risk of vector toxicity and ocular inflammation. In this study, we subretinally administered a low dose AAV2/4-EF1α-*RS1* gene therapy to *Rs1*-KO mice, a model for XLRS. To account for the potential effect of the subretinal injection procedure, a sham control group receiving a hypertonic buffer, similar in osmolarity to the AAV storage buffer, was included.

We hypothesized that subretinal injections of low dose AAV2/4-EF1α-*RS1* could correct the retinal phenotype in the *Rs1*-KO mouse detectable by electroretinography (ERG), optical coherence tomography (OCT), and a visually guided swim assay (VGSA). However, our experiments revealed an unexpected therapeutic effect of the buffer, which led to further exploration of this effect.

## 2 Methods

### 2.1 Study Design

Natural history, as well as gene therapy efficacy and effect of subretinal buffer injections, were evaluated in a mouse model of Juvenile X-Linked Retinoschisis (*Rs1*-KO). The natural history of this *Rs1*-KO mouse model has been reported, including the variable expressivity of the phenotype (12). We examined treatment-naïve eyes from postnatal day (P) 15 to 12 months of age. OCT images were analyzed for both outer nuclear layer (ONL) thickness and cyst severity.

To study the effect of gene therapy, eyes treated with a subretinal injection of gene therapy vector (AAV2/4-EF1α-*RS1*) (n=10) were compared to hypertonic buffer-injected eyes (n=16) and treatment-naïve eyes (n=27). Injections alternated between the left and right eyes to avoid systematic bias.

The selection of an AAV2/4 vector was based on previously demonstrated tropism for all layers of the retina, including the photoreceptor and bipolar cell layers, the primary secreters of retinoschisin (13).

Injections were performed between P24 and P31. Outcome measures include ERG, OCT, VGSA, and immunohistochemistry. ERG was performed at 1-, 2-, 3-, and 5-months post injection (MPI) to assess retinal function. OCT was performed at 2- and 3-weeks post-injection (WPI), then 1-, 2-, 3-, and 5 MPI to assess cyst severity and retinal structure. A visually guided swim assay was performed at 5-6 MPI to assess functional vision. The eyes were collected and fixed at 6 MPI for immunohistochemistry. For the purpose of ERG and OCT experiments, one eye is considered a biological data point. For the purpose of the visually guided swim assay, each data point is representative of the average time-to-platform for a randomly selected platform position. Mice were treated in one eye with either vector or hypertonic buffer. Mice treated with gene therapy in one eye and hypertonic buffer in the fellow eye were excluded from the VGSA. Both male (*Rs1* hemizygous mutants) and female mice (*Rs1* homozygous mutants) were used in this study.

To investigate the effect of the buffer, the subretinal injection of the hypertonic buffer was compared to the subretinal injection of an isotonic buffer. Endpoints included ERG and OCT, and the same timelines as stated above were used in this group.

### 2.2 Animal Husbandry and Ethics Statement

This study was performed in accordance with the recommendations set forth by the National Institute of Health in the Guide for the Care and Use of Laboratory Animals. Animals were handled in accordance with the approved Institutional Animal Care and Use Committee (IACUC) protocol #1041421 at the University of Iowa. The *Rs1*-KO C57Bl/6J mouse model was kindly supplied by Paul Sieving, M.D., Ph.D. at the National Eye Institute. Generation of the *Rs1*-KO mouse model was previously described (14). This mouse model contains a deletion of exon 1 and a 1630 bp fragment of intron 1 on the *Rs1* gene (14).

Animals were housed according to the IACUC recommendations. Humane endpoints were observed, and the methods of euthanasia used were carbon dioxide inhalation followed by cervical dislocation.

### 2.3 Statistical Analysis

Analysis was performed using GraphPad PRISM 9.0 (GraphPad Software, Inc, San Diego, California, USA). Two-way ANOVA was performed for ERG and OCT metrics and followed by multiple comparisons (Tukey’s test, non-parametric). One-way ANOVA was performed for VGSA and followed by multiple comparisons (Tukey’s test, non-parametric). The student’s t-test was used in the analysis of cone density. Simple linear regression was used for correlation of peak cyst area vs 5 MPI ERG response.

### 2.4 Genotyping Information

Genotyping was performed using Taq Polymerase (New England biolabs, Ipswich, MA) using the primers listed in Table 1. The PLA2 primer was previously described (14). A 10 microliters (μL) reaction volume was used containing 2 μL of DNA (approximately 25 ug/μL), 3.7 μL of ultrapure water, 1 μL of 10X Buffer/Mg++, 1 μL of 20 mM primer mix (Table 1), 0.2 μL of 50 mM dNTPs, 0.1 μL of rTaq, and 2 μL of 5X Betaine. Cycling conditions are as follows; initial denaturing at 94°C for 30 seconds, annealing at 57°C for 30 seconds and extension at 72°C for 30 seconds, with a final extension step at 72°C for 4 minutes. This produces a 516-base pair (bp) wild-type band and a 300 bp knockout band.

**Table 1:**
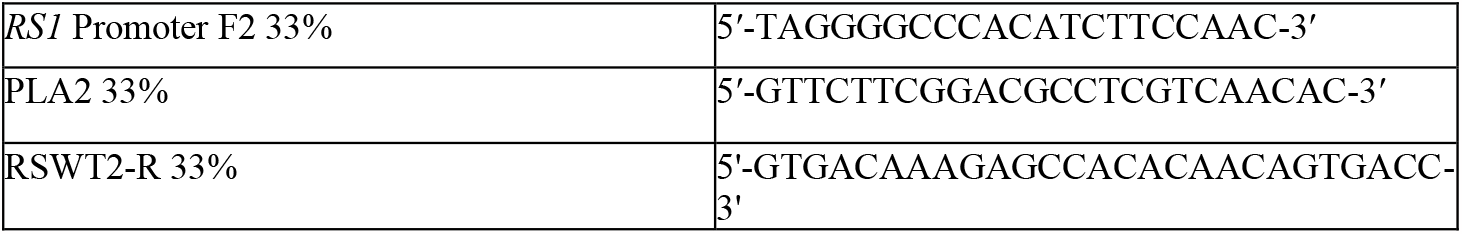
Primers.

### 2.5 AAV Packaging

The human *RS1* gene was cloned into a shuttle plasmid pFBAAV and provided by the Viral Vector Core at the University of Iowa. The bacterial backbone is from Invitrogen’s pFastBac system. The shuttle plasmid consists of the pFBAAV backbone, an EF1α promoter, and a bovine growth hormone polyadenylation signal. The plasmid was packaged into an AAV2/4 capsid by the Budd Tucker Laboratory at the University of Iowa. The AAV was formulated at 2 × 10^9^ viral genomes (vg)/mL titer, and the composition of the storage buffer is listed in Table 2. Titering was performed using digital droplet PCR.

**Table 2:** Buffer solutions.

|  |  |
| --- | --- |
| AAV storage buffer | • 180 mM NaCl<br>• 10 mM Na<sub>3</sub>PO<sub>4</sub><br>• UltraPure water<br>• pH 7.2-7.4 |
| hypertonic injection buffer | • 0.001% Gibco Pluronic F-68<br>• Gibco dPBS (-/-(Catalog #14190144) [Contains 155.2 mM NaCl]<br>• Additional 24.8 mM NaCl (for 180 mM NaCl total)<br>• pH 7.4 |
| isotonic injection buffer | • 0.001% Gibco Pluronic F-68<br>• Gibco dPBS (-/-(Catalog #14190144) [Contains 155.2 mM NaCl]<br>• pH 7.4 |

### 2.6 Subretinal Injection

Mice were anesthetized using a ketamine/xylazine mixture (87.5 mg/kg ketamine, 12.5 mg/kg xylazine). A tropicamide ophthalmic solution of 1% was applied three minutes prior to injection to dilate the eyes. Proparacaine was applied as a topical anesthetic and 10% povidone-iodine was applied as a topical antiseptic. A partial thickness puncture was made just posterior to the limbus through the sclera with a 30-G sharp needle, then a 33-G blunt Hamilton needle was inserted and positioned between the RPE and the retina. Two μL of either gene therapy vector at a concentration of 2 × 10^9^ vg/mL, a hypertonic buffer, or an isotonic buffer was injected into the subretinal space (formulation available in Table 2). The injection of a solution subretinally creates a separation of the retina and the RPE known as a subretinal bleb. This bleb was assessed by the injector as seen through the operating microscope and in the rare cases with no visible bleb or a large vitreous hemorrhage, the mouse was excluded. Additionally, mice with retinal detachments that persisted until the first OCT time point were excluded. The mouse was then given topical ophthalmic ointment consisting of neomycin, polymyxin B sulfates, bacitracin zinc, and hydrocortisone. Mice were injected with atipamezole to aid in recovery from the anesthetic.

### 2.7 Electroretinogram

Dark adaptation occurred overnight. Mice were anesthetized using a ketamine/xylazine mixture (87.5 mg/kg ketamine, 12.5 mg/kg xylazine). A tropicamide ophthalmic solution of 1% was applied to the eyes for pupil dilation. A 2.5% Hypromellose solution (Akron, Lake Forest, Illinois) was applied to maintain corneal hydration during testing. Testing was performed on the Diagnosys Celeris (Diagnosys, Lowell, MA). Electrodes were placed on the cornea of bilateral eyes, and impedance was kept below 10 kΩ. Eyes were exposed to various light intensities in a modified ISCEV protocol which includes a dark-adapted 0.01 cd•s/m^2^, dark-adapted 3.0 cd•s/m^2^, light-adapted 3.0 cd•s/m^2^, and a light-adapted 5 Hz flicker of 3.0 cd•s/m^2^ which is more reproducible in mice than the standard light-adapted 30 Hz flicker in humans (15). To evaluate rod electrical function in treated and untreated *Rs1*-KO mice, animals were dark-adapted (DA) overnight to isolate the rod response, and eyes were subjected to 15 flashes of 0.01 cd•s/m^2^ light (0.01 dim flash) then 15 flashes of 3.0 cd•s/m^2^ light (3.0 standard combined response (SCR)). To evaluate cone electrical function in treated and untreated *Rs1*-KO mice, eyes were light-adapted (LA) for 10 minutes to bleach the responses from rods and subjected to 15 flashes of 3.0 cd•s/m^2^ light (3.0 flash) followed by 20 flashes of a 5 Hz flickering light of 3.0 cd•s/m^2^ (5 Hz Flicker).

### 2.8 Optical Coherence Tomography

Mice were anesthetized using a ketamine/xylazine mixture (87.5mg/kg ketamine, 12.5mg/kg xylazine), and Tropicamide ophthalmic solution 1% was applied for pupil dilation. 1% carboxymethylcellulose was applied to maintain corneal hydration during testing. Volumetric scans were taken centering on the optic nerve along with nasal and temporal scans. Nasal and temporal scans involved moving the optic nerve peripherally, as previously described (16). Quantification of cyst severity was performed using two different methods. The first method consisted of a modified scoring protocol from previously described literature (17). Briefly, measurements of cyst height were taken at four points equidistant, 500 μm from the optic nerve, two on the superior-inferior line bisecting the optic nerve, and two on the nasal-temporal line bisecting the optic nerve. Measurements were then translated to a scoring scale as previously described, in which a score of 1 was assigned to zero cyst height and a score of 6 was assigned to cyst height >100 μm. The score was averaged across all four points (16). The second method consists of the manual segmentation of cyst area.

Briefly, OCT images were analyzed for cyst area using the Image J software. Center scans, those which best bisect the optic nerve, were quantified by manually segmenting individual cyst cavities and recording the total area of cysts in the scan. Similar quantifications were performed on nasal-temporal slices located approximately 500 μm superior and inferior of the center scan, and area was reported as a sum of these measurements.

### 2.9 Visually Guided Swim Assay

AAV2/4-EF1α-*RS1* vector-treated mice and mice injected with the hypertonic buffer underwent a VGSA performed in light and dark conditions to assess functional vision at approximately 5-6 months of age compared to treatment-naïve controls. The VGSA has been described in detail elsewhere (18). In brief, the mice undergo four training days in the light, four testing days in the light, two training days in the dark, and four testing days in the dark, and the performance of each mouse to 5 different platform locations were measured during each training or testing day. For each trial, the mice were placed into a plastic pool and were expected to find a platform. Five of the eight possible platform positions are randomly assigned each day with all platforms being used approximately the same number of times. A maximum time to platform (TTP) was set at 60 seconds for each trial to prevent fatigue and undue stress. After the mouse had completed the trial or after 60 seconds of searching, the mouse was removed from the pool, dried, and placed in its cage until the next trial. Mice were excluded in rare cases where they could not be motivated to participate and search for the platform. Data from all trials was excluded if a mouse had a swim time greater than one standard deviation from the mean of the group and consistent floating occurred, such as more than three episodes of floating per trial or more than three corrections per trial. Corrections for floating include snapping fingers or touching the mouse’s tail.

### 2.10 Immunohistochemistry

Eyes were enucleated and a small puncture was made through the cornea with a 27-G needle. Eyes were placed in 4% paraformaldehyde for 12 hours and transferred to PBS for 24 hours. After fixation, eyes were embedded in Tissue-Tek O.C.T. compound (VWR, Batavia, IL) and frozen in a bath of 2-methylbutane cooled with liquid nitrogen. Eyes were sectioned on the superior-inferior axis at a thickness of 10 μm using a Cryostat microtome and stored at -80°C for future use.

For immunohistochemistry, sections were permeabilized using 0.3% Triton X-100 in PBS for 10 minutes at room temperature, blocked with a blocking buffer containing 5% BSA, 5% normal goat serum, and 0.05% Triton X-100 in PBS for 1 hour at room temperature, and then incubated with primary antibodies in a dilution buffer containing 5% BSA, 1% normal goat serum, and 0.05% Triton X-100 in PBS at 4 °C overnight. The following day, samples were washed in PBS three times before incubation with secondary antibodies or streptavidin Alexa Flour-568 conjugate at room temperature for 1 hour. After washing in PBS an additional three times, samples were mounted with VectaShield mounting medium with DAPI (Vector Laboratories, Burlingame, CA). Images were taken using a fluorescence microscope and color channels were overlayed in Image J. No contrast enhancements or brightness levels were altered after acquisition.

Antibodies used for immunohistochemistry were as follows: anti-CtBP2/RIBEYE antibody (BD Transduction Laboratories #612044; 1:200 dilution); biotinylated-peanut agglutinin (biotinylated-PNA) (Vector Laboratories #B-1075; 1:500 dilution).

Images were taken directly superior and inferior of the optic nerve head and were gathered at 40x magnification. Three serial images were acquired on either side of the optic nerve leading to six total images gathered per eye. These images were then independently quantified by three individuals masked to treatment groups and the number of cones per image was averaged. The quantifications were averaged and divided by the image width to produce the reported average cones per 100 μm.

## 3 Results

### 3.1 Natural history of disease in the *Rs1*-KO mouse compared to WT BL/6 mouse

Cyst severity was quantified over time in the *Rs1*-KO mouse using a modified cyst severity scoring system, as previously described by Bush et al. (17). Figure 1 demonstrates typical OCT findings in the *Rs1*-KO mouse at P15, and 2-, 3-, and 6-months of age, compared to a WT mouse (Figure 1A). At all-time points, cysts, or schisis, can be observed in the *Rs1*-KO mouse retina. This compares to the organized, laminar retinal architecture of the WT mouse. Cysts are apparent as early as P15 with an average severity score of 2.75 (Figure 1C). Cysts appear the most severe at 2- to 4-month-old, hitting a local maxima at 3 months. Thereafter, severity reduced with maturity.

**Figure 1.**
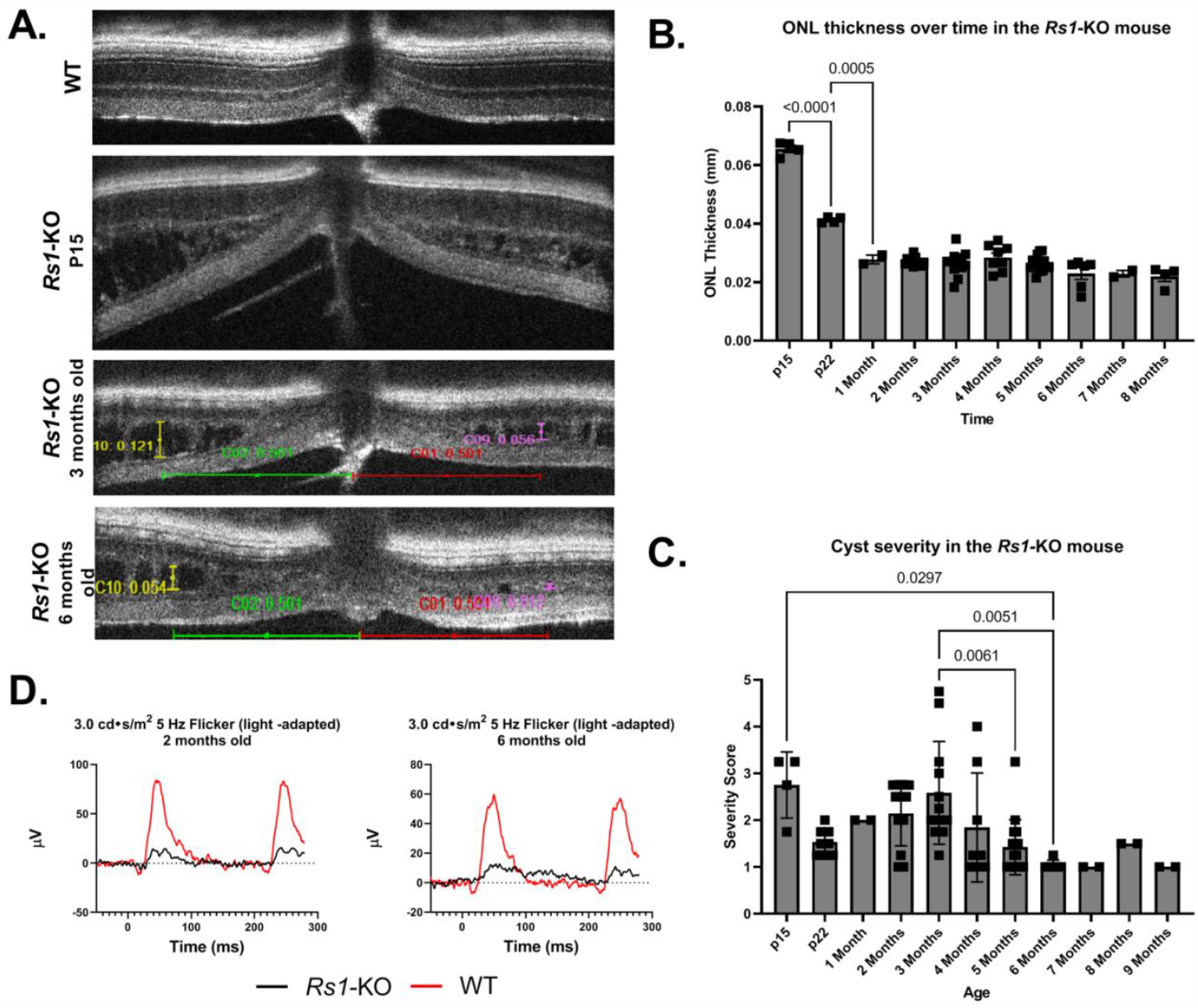
(A-D): Natural history of disease in the *Rs1*-KO mouse compared to WT BL/6 mouse. OCT images were collected from p15 to 9 months of age in *Rs1*-KO mice, and ONL thickness (B) and cyst severity (C) were measured at each age. Measurements of cyst height were taken at four points equidistant, 500 um, from the optic nerve, and the averaged values for each eye were reported. For cyst severity, cyst height was measured and translated to a scoring scale of 1 to 6, with 1 representing a cyst height of 0. Representative OCT images demonstrate retinoschisis in *Rs1*-KO mice at P15, 3-, and 6 months old, compared to a WT mouse without retinoschisis (A). ERGs of *Rs1*-KO mice and WT mice were collected at 2 and 6 months of age. As early as 2 months post-injection, *Rs1*-KO mice show reduced ERG B wave amplitudes and delayed latency of the b wave peak, compared to the robust WT waveforms (D).

In untreated *Rs1*-KO mice, significant thinning of the ONL occurred from P15 to 1 month of age (Figure 1B, *p<0*.*0001*). After 1 month, the ONL thickness in *Rs1*-KO mice did not significantly change over time, indicating that the majority of the photoreceptor cell loss occurred shortly after birth and during eye maturation.

ERGs investigate the electrical response of the retina *in vivo* to understand how the disease affects signal transmission in light and dark conditions. ERGs of *Rs1*-KO mice demonstrated consistently reduced function of rod-dependent and cone-dependent visual pathways compared to WT. Figure 1 demonstrates typical ERG waveforms for dark-adapted (3.0 SCR) and light-adapted (5 Hz flicker) metrics (Figure 1D).

In summary, the *Rs1*-KO mouse model closely resembles the clinical course in humans by demonstrating cyst formation, significant ONL thinning, and diminished electrical activity of retinal photoreceptors and bipolar cells observable on ERG. In our hands, this model has a phenotype similar to that described by Sieving et al. (12).

By treating at 1 month of age, we aimed to determine if we could mitigate the formation of severe cysts and subsequently improve the electrical signaling of retinal cells in adult mice after the initial ONL thinning phase. Results from vector-treated and hypertonic buffer-injected eyes were compared to those of treatment naïve eyes (untreated).

### 3.2 Cyst severity is significantly reduced in vector-treated and hypertonic buffer-injected eyes at 1 MPI

Using a modified cyst severity scoring system, vector-treated *Rs1*-KO eyes (n=10) demonstrated significantly less cyst severity than untreated eyes (*p=0*.*002*; Figure 2B) at 1 MPI. Hypertonic buffer-injected eyes also showed a similar reduction in cyst severity to compared to untreated eyes at 1 MPI (*p<0*.*0001*, Figure 2B). The observed reduction in cyst severity occurred at 1 MPI at which point the mice are 2 months of age. As previously described, peak cyst formation in *Rs1*-KO mice begins around 2 months of age, suggesting injection of the vector or hypertonic buffer leads to reduction in cysts at a significant time in cyst development.

**Figure 2.**
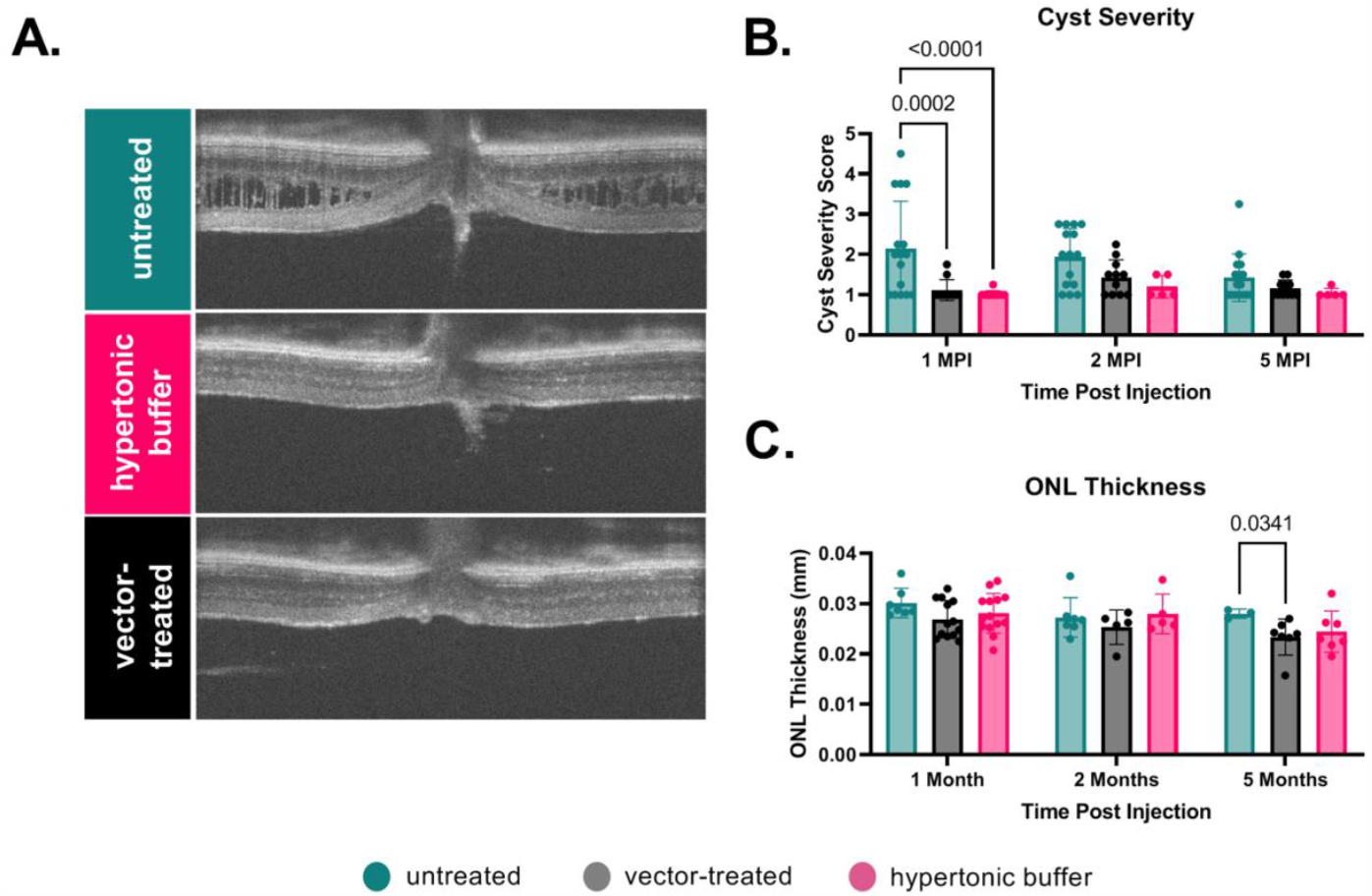
(A-B): Cyst severity is significantly reduced in vector-treated and hypertonic buffer-injected eyes at 1 MPI. OCT images of vector-treated, hypertonic buffer-injected, and untreated eyes were collected at 1, 2, and 5 MPI. Measurements of cyst severity (B) and ONL thickness (C) were taken at four points equidistant, 500 um, from the optic nerve, and each datapoint is the averaged value of an eye. For cyst severity, cyst height was translated to a modified scoring scale of 1 to 6, and averaged for that eye. Both vector-treated (*p=0*.*002*) and hypertonic buffer-injected eyes (*p<0*.*0001*) had significantly less cysts than untreated eyes at 1 MPI, and continued to have few to no cysts over time. Representative OCT findings at 1 MPI (A).

At 2 and 5 MPI no discernible distinction in cyst severity was evident among untreated, vector-treated, or hypertonic buffer-injected mice. The attenuation of significance observed in these comparisons can be ascribed to the previously established reduction in cyst severity observed in untreated *Rs1*-KO mice, per the natural disease progression, rather than an increase in cysts within the injection groups (12). ONL thickness remained stable from 1 to 5 MPI in all cohorts, and perceived significance at 5 MPI is likely an anomalous data (Figure 2C).

Cysts spatially separate bipolar cells and photoreceptors, and it is hypothesized that the lower b-wave amplitudes and/or electronegative ERG phenotype associated with XLRS is a consequence of this spatial separation. To examine the potential influence of reduced cysts on retinal electrical signaling, we conducted ERGs under light-adapted and dark-adapted conditions.

### 3.3 Hypertonic buffer outperforms untreated and vector-treated eyes in light-adapted ERGs, but not in dark-adapted ERGs

Vector-treated *Rs1*-KO eyes (n=10) or hypertonic buffer-injected *Rs1*-KO eyes (n=16) were compared to untreated *Rs1*-KO eyes.

#### 3.3.1 Light-Adapted

The retinal cone pathway was measured using two light-adapted ERG assays; the light adapted 5 Hz flicker and 3.0 flash. At 1 MPI, the vector-treated eyes and hypertonic buffer-injected eyes demonstrated superior cone function to untreated eyes in the 5 Hz flicker assay (v*ector-treated; p=0*.*0003; hypertonic buffer p<0*.*0001;* Figure 3A). This effect persisted to 5 MPI (v*ector-treated: p=0*.*0255; hypertonic buffer: p=0*.*0018;* Figure 3A). Although both injection groups were superior to untreated controls, hypertonic buffer-injected eyes had significant improvement over vector-treated eyes at the 5 MPI time point (*p=0*.*0235*), suggesting a more robust cone rescue occurred after subretinal injection of a hypertonic buffer compared to the low-dose AAV2/4-EF1α-*RS1* vector.

**Figure 3.**
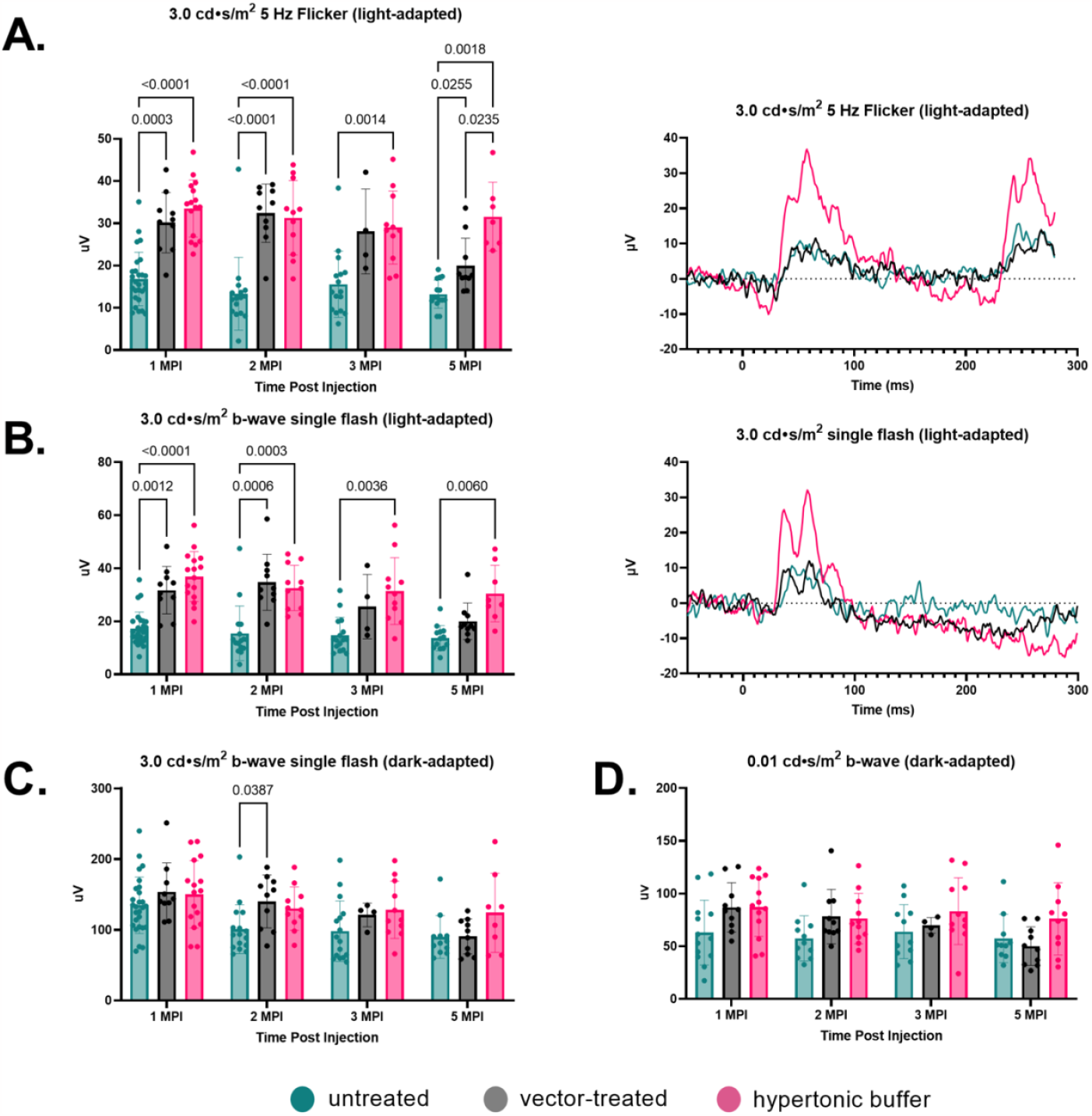
(A-D): Hypertonic buffer outperforms untreated and vector-treated eyes in light-adapted ERGs, but not in dark-adapted ERGs. Cone function was measured using two separate light-adapted ERG assays - the 5 Hz Flicker and 3.0 cd•s/m^2^ bright flash. In response to the 5 Hz flicker stimulus, vector-treated eyes and hypertonic buffer-injected eyes had higher amplitudes than untreated eyes at 1, 2, 3, and 5 MPI (5 MPI: *vector-treated: p=0*.*0255; hypertonic buffer: p=0*.*018*) (A). Hypertonic buffer-injected eyes were also superior over vector-treated eyes at the 5 MPI time point (*p=0*.*0235*) (A). Under the 3.0 cd•s/m^2^ bright flash assay, an additional measurement of cone-pathway function, vector-treated eyes demonstrated superior cone function to untreated eyes till 2 MPI (*p=0*.*0006*), whereas the hypertonic buffer-injected eyes were superior till 5 MPI (*p=0*.*006*) (B). (In summary, hypertonic buffer-injected eyes had a more robust and long-term improvement in cone-dominant pathways.) Representative waveforms demonstrate these findings at 5 MPI (A-B). Combined rod-cone function was measured by the 3.0 cd•s/m^2^ bright flash under dark-adapted conditions (C). At 1, 3, and 5-MPI, there was no difference between vector-treated, hypertonic buffer-injected, or untreated eyes. At 2 MPI, vector-treated eyes did demonstrate superior function over untreated eyes, but this effect did not persist. Isolated rod function was measured using 0.01 cd•s/m^2^ dim flash after dark adaptation (D). There was no persistent significant difference between hypertonic buffer-injected, isotonic buffer-injected, or untreated eyes in rod dominant ERG settings, suggesting no or minimal rod rescue following injections.

In the 3.0 flash, vector-treated eyes demonstrated superior cone function to untreated eyes till 2 MPI (v*ector-treated: p=0*.*0006; hypertonic buffer: p=0*.*0003;* Figure 3B). However, hypertonic buffer-injected eyes had a more robust and long-term improvement in function, demonstrating superior cone function over untreated eyes out to 5 MPI (*p=0*.*006*). The robust and long-term improvement in cone functioning observed in the hypertonic buffer group was a surprising finding and indicates that the subretinal injection of buffer alone enabled long-term benefits in retinal cone function. Representative waveforms demonstrate these findings at 5 MPI (Figure 3, A-B).

#### 3.3.2 Dark-Adapted

To measure the electrical function of the eye in dark conditions, isolating the rod response, the 3.0 SCR and 0.01 dim flash ERG assays were utilized. We found no significance between the untreated, vector-treated, or hypertonic buffer-injected eyes at 1 MPI in either 3.0 SCR *(vector-treated: p=0*.*433; hypertonic buffer: p=0*.*531*, Figure 3C), or 0.01 dim flash (v*ector-treated: p=0*.*28, hypertonic buffer: p=0*.*158*, Figure 3D). No significant effect was observed up to 5 MPI in the dark, suggesting that subretinal injections of low-dose AAV2/4-EF1α-*RS1* vector or buffer had minimal impact on rod-dependent retinal function.

Following the observed improvements in cone specific electrical pathways, we investigated whether these improvements were associated with differences in functional vision.

### 3.4 Vector-treated and hypertonic buffer-injected mice perform faster in a light-adapted swim assay than untreated *Rs1*-KO mice

To assess functional vision, mice subretinally injected with the vector or hypertonic buffer were subjected to a VGSA and compared to untreated *Rs1*-KO mice. Mice swam at 5-6 months of age, in light- and dark-adapted environments, and were timed until they found a platform in the pool. In light-adapted swimming conditions, we observed that vector-treated mice (2.92 seconds) were significantly faster at finding the platform than untreated mice (5.06 seconds) (*p=0*.*0002*) (Figure 4A). The hypertonic buffer-injected mice (2.51 seconds) were also significantly faster than untreated mice when swimming in a lighted environment (*p<0*.*0001*). These results confirm that the subretinal injection of vector or buffer alone provided a benefit in functional vision in the light, which correlates with the apparent reduction of cysts, and improved cone pathway ERG signaling. There was no observable significance between vector-treated, hypertonic buffer-injected, and untreated mice in a dark-adapted swim environment (Figure 4B).

**Figure 4.**
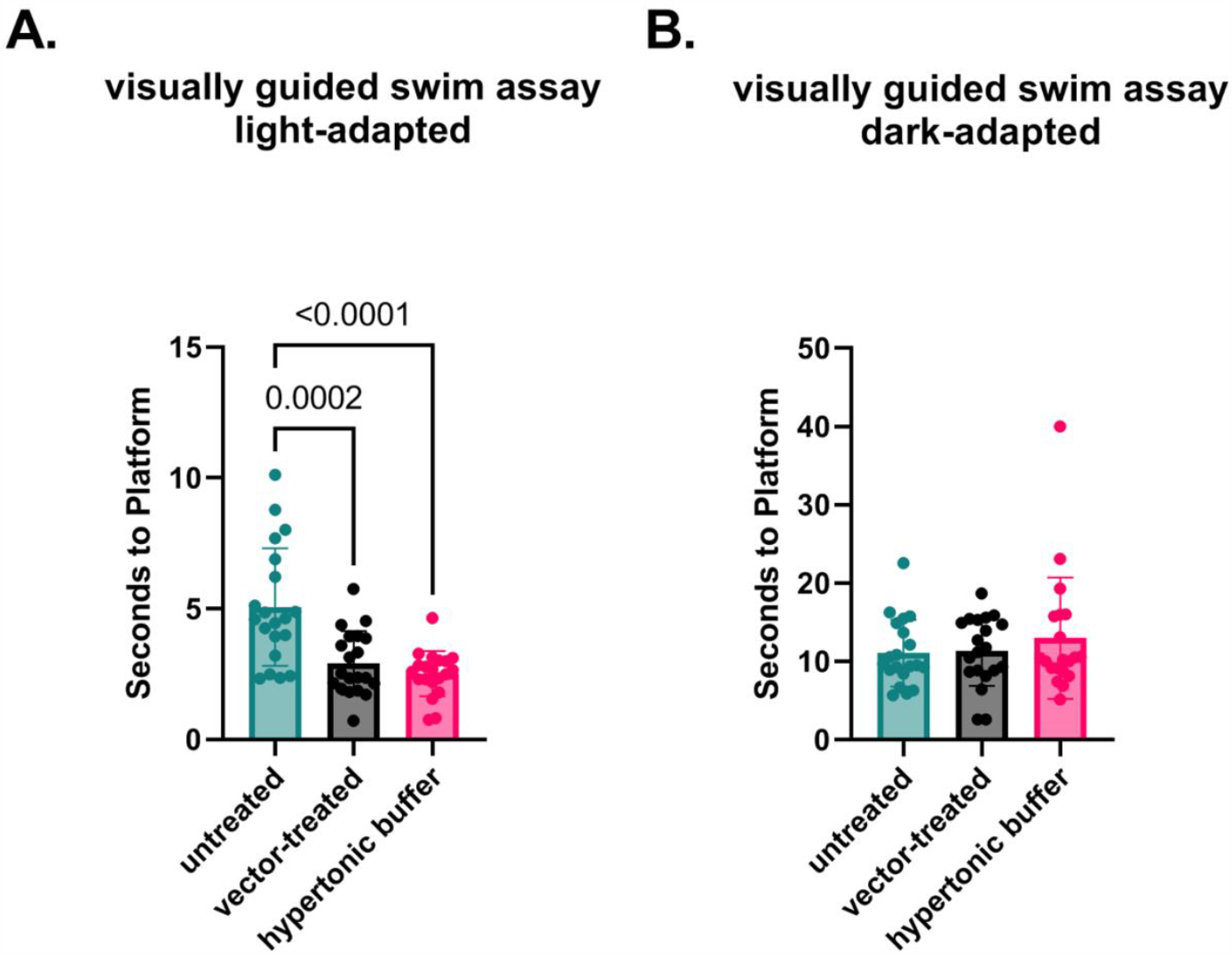
(A-B): Vector-treated and hypertonic buffer-injected mice perform faster in a light-adapted swim maze than untreated *Rs1*-KO mice. Mice were placed in a swim maze and testing was performed under dark or light adaptation conditions. Swim time to a platform was recorded. Each data point is representative of the average time-to-platform for a randomly selected platform position. In a light-adapted environment, vector-treated and hypertonic buffer-injected mice are significantly faster at finding swim platforms than untreated *Rs1*-KO mice (*vector-treated: p=0*.*0002; hypertonic buffer: p<0*.*0001*), suggesting improvement in light-adapted visual acuity (A). In a dark-adapted environment, there is no significant difference in time-to-platform between vector-treated, hypertonic buffer-injected, and untreated eyes, suggesting no effect on visual acuity in the dark (B).

Together, long-term ERG data and functional vision assay suggest that subretinal injections of a hypertonic buffer solution alone led to reductions in cyst severity and cone-specific improvements in retinal function as well as functional vision. The improvements observed in buffer-treated eyes were equivalent or sometimes superior to improvements observed in vector-treated eyes.

### 3.5 Investigation of the effect of tonicity on cone rescue

Our data suggests that the injection of hypertonic buffer alone leads to significant improvements in cone-specific ERG amplitudes, light-adapted visual function, and retinal architecture in *Rs1*-KO mice. Previous studies have demonstrated a temporary reduction in existing retinal cavities following the subretinal injection of a balanced salt solution; however, the long-term effects on visual function and electrical signaling of the retina has not been studied (19).

Additionally, cystoid retinal conditions in XLRS patients can be modified by the application of diuretic ophthalmic solutions such as brinzolamide and acetazolamide, but the connection between fluid reabsorption and cyst resolution is not well understood. Previous literature has suggested that the retinoschisin protein physically interacts with Na/K-ATPase and may act to mediate Na/K-ATPase activity, affecting signaling and ion gradients within the retina (20, 21). We noted that the buffer solution, used both to store the AAV vector and used in the initial buffer injections, contained a high salt content achieving a greater than physiologic osmolarity. Therefore, we explored the idea that the tonicity of the injected buffer played a role in promoting cyst resolution and improving retinal signaling. To investigate this hypothesis, we repeated experiments with a subretinal injection of an isotonic buffer. The isotonic buffer has a similar composition to the hypertonic buffer except possessing a lower concentration of NaCl. The endpoints of these experimental groups also include ERG to measure retinal electrical function and OCT to measure retinal structure and cyst severity.

The results for these tests are as follows.

### 3.6 Cyst area is reduced in hypertonic buffer and isotonic buffer-injected eyes, with greater reduction observed in the hypertonic buffer group

To determine if structural differences exist when modulating the tonicity of the injection buffer, OCT was used to image the retinal layers *in vivo*. Using ImageJ software, images were then assessed for cyst severity using manual segmentation and measurement of visible cyst cavities reported as total cyst area. Outer nuclear layer (ONL) thickness measurements were taken to assess photoreceptor cell survival.

We observed less cyst severity in the hypertonic buffer and isotonic buffer-injected eyes compared to untreated eyes at 3 weeks post-injection (WPI) (*hypertonic buffer: p=0*.*0033, isotonic buffer: p=0*.*0431*, Figure 5A). By 1 MPI, hypertonic and isotonic buffer-treated eyes continued to have less severe cysts than untreated eyes (h*ypertonic buffer: p<0*.*0001, isotonic buffer: p<0*.*0001*). This effect persisted until 2 MPI for the hypertonic buffer-injected eyes (*p=0*.*0380*). In summary, hypertonic buffer-injected eyes demonstrated an early reduction in cyst severity, originating as early as 3 weeks PI and this effect persisted to 2 MPI. Additionally, hypertonic buffer-injected eyes had a reduction of severity to a slightly greater magnitude than isotonic buffer-injected eyes at 3 WPI and 1 MPI. Cyst area was decreased across all groups by 5 MPI.

**Figure 5.**
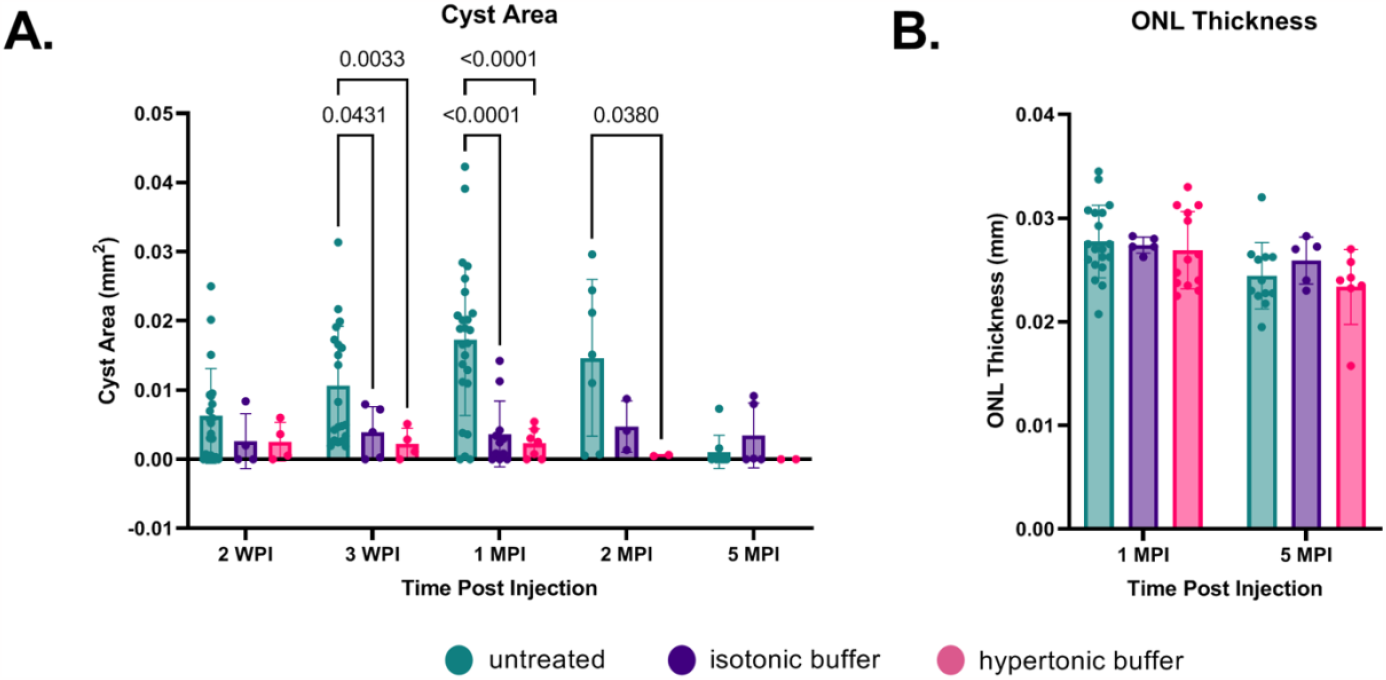
(A-B): Cyst area is reduced in hypertonic buffer and isotonic buffer-injected eyes, with greater reduction observed in the hypertonic buffer group. OCT images were collected at 2- and 3-weeks post injection (WPI), 1 MPI, and 2 MPI. Cyst area quantification and outer nuclear layer (ONL) thickness measurements were taken to assess photoreceptor cell survival. Each data point is representative of one eye. Starting at 3 WPI, cyst area is significantly reduced in hypertonic buffer and isotonic buffer-injected eyes compared to untreated eyes (3 WPI: *hypertonic buffer: p=0*.*0033; isotonic buffer: p= 0*.*0431*). For the hypertonic-buffer injected eyes, this effect persisted to 2 MPI (2 MPI: *hypertonic buffer: p=0*.*0380)* (A). Cyst area is decreased across all groups at 5 MPI (A). ONL thickness remains stable to 5 MPI and does not significantly differ between injection groups and untreated controls (B).

Though cysts were decreased at 1 MPI, ONL thickness was not different across time points between hypertonic buffer-injected, isotonic buffer-injected, and untreated eyes (h*ypertonic buffer: p=0*.*8995, isotonic buffer: p=0*.*9916, hypertonic buffer vs isotonic buffer: p=0*.*9921*, Figure 5B).

Having observed a notable decrease in cysts among mice subjected to a subretinal injection of hypertonic buffer, and to a lesser effect isotonic buffer, we investigated whether cyst reduction paralleled improvements in electrical functioning of the retina.

### 3.7 Hypertonic buffer-injected eyes have a more robust cone rescue on light-adapted ERG over untreated and isotonic buffer-injected eyes

*Rs1*-KO eyes were subretinally injected with hypertonic buffer solution (n=16) or isotonic buffer solution (n=7) and compared to untreated eyes (n=27).

#### 3.7.1 Light-Adapted

Following an observed reduction in retinal cysts after subretinal injection of a hypertonic buffer, we investigated whether the improvement in cysts paralleled a similar improvement in electrical function of the retina.

Cone function was measured using two separate light-adapted ERG assays; the light adapted 3.0 flash and 5 Hz flicker. Under the 3.0 flash following light adaptation, both isotonic buffer and hypertonic buffer-injected eyes displayed superior cone function over untreated eyes, persisting from 1 MPI *(hypertonic buffer: p<0*.*0001, isotonic buffer: p=0*.*0192*, Figure 6A) to 5 MPI *(hypertonic buffer: p=0*.*0060, isotonic buffer: p=0*.*0089;* Figure 6A). Although both buffer groups demonstrated an improvement in cone pathway signaling over untreated eyes, hypertonic buffer-injected eyes exhibited a more pronounced improvement.

**Figure 6.**
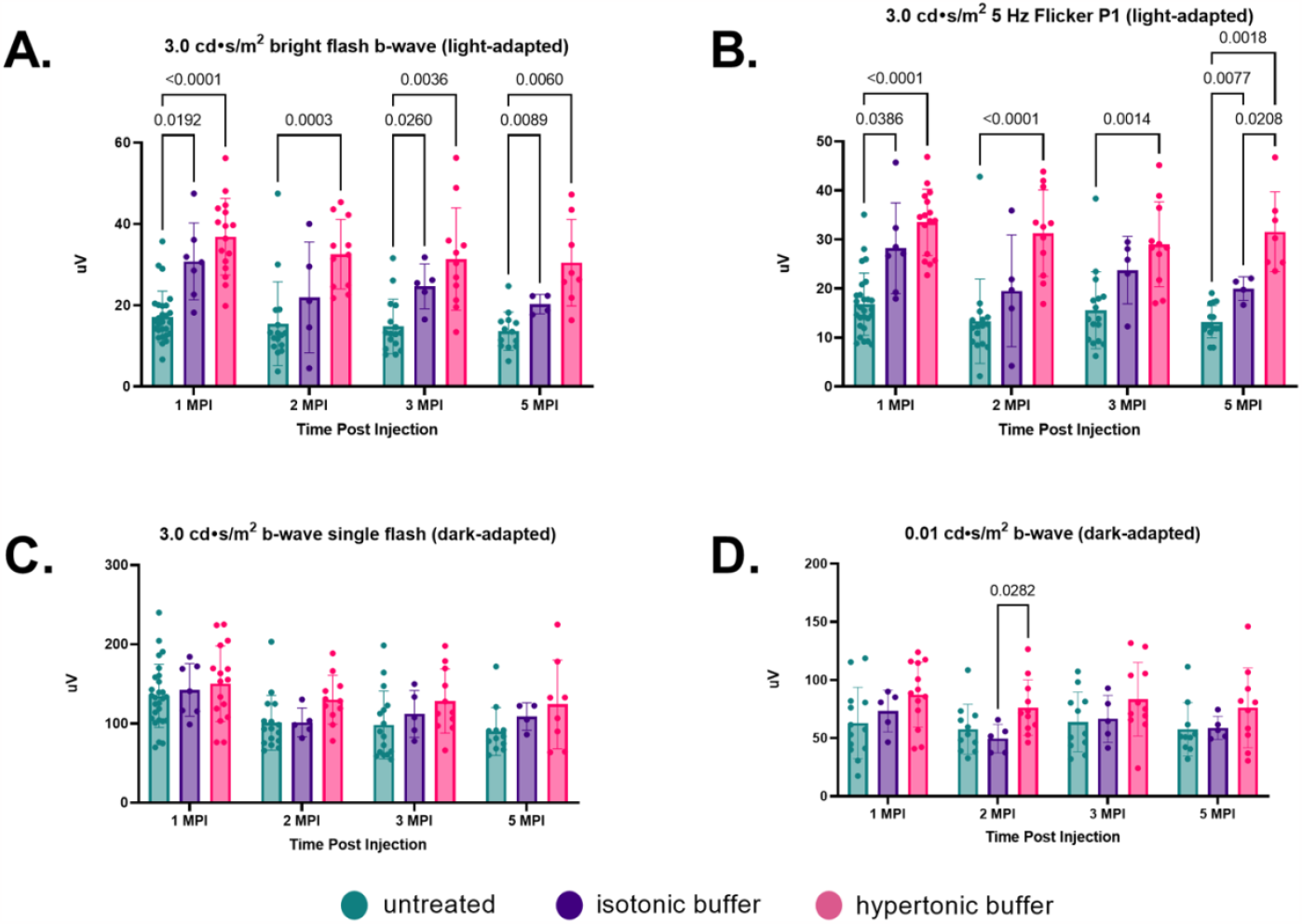
(A-D): Hypertonic buffer-injected eyes have a more robust cone rescue on light-adapted ERG over untreated and isotonic buffer-injected eyes. To determine if tonicity had a potential effect on improving electrical functioning of the retina, an isotonic buffer was subretinally introduced into the eyes of *Rs1*-KO mice. Cone function was measured using two separate light-adapted ERG assays - the 5 Hz Flicker and 3.0 cd•s/m^2^ bright flash. Each data point is representative of one eye. In response to the 3.0 cd•s/m^2^ bright flash, both isotonic buffer and hypertonic buffer-injected eyes displayed superior cone pathway functioning over untreated eyes, persisting to 5 MPI (5 MPI: *hypertonic buffer: p=0*.*0060; isotonic buffer: p=0*.*0089*) (A). However, hypertonic buffer-injected eyes exhibited a more pronounced improvement over untreated controls compared to isotonic buffer-injected eyes. Under the 5 Hz flicker assay, isotonic buffer and hypertonic buffer-injected show improved functioning over untreated eyes to 5 MPI (5 MPI: *hypertonic buffer: p=0*.*0018; isotonic buffer: p=0*.*0077*). However, hypertonic buffer-injected eyes were superior to isotonic buffer-injected eyes at 5 MPI (*p=0*.*0208*). (In summary, hypertonic buffer-injected eyes had a more robust and long-term improvement in cone-dominant pathways). The combined rod-cone function was measured by the 3.0 cd•s/m^2^ bright flash under dark-adapted conditions (C). Isolated Rod function was measured using 0.01 cd•s/m^2^ dim flash after dark adaptation (D). There was no persistent significant difference between hypertonic buffer-injected, isotonic buffer-injected, or untreated eyes in rod dominant ERG settings, suggesting no or minimal rod rescue following injections.

In the 5 Hz flicker protocol, isotonic buffer-injected eyes showed some improved functioning over untreated eyes from 1 MPI (*isotonic buffer: p=0*.*0386*, Figure 6B) to 5 MPI *(isotonic: p=0*.*0077*, Figure 6B). Though there are observable improvements in cone function in the isotonic buffer-injected eyes, the magnitude and significance of these improvements are less than the improvements observed in the hypertonic buffer-injected eyes. Hypertonic buffer-injected eyes demonstrated a more robust improvement in cone function and were superior to isotonic buffer-injected and untreated eyes up to 5 MPI *(hypertonic buffer vs untreated: p=0*.*0018; hypertonic buffer vs isotonic buffer: p=0*.*0208*, Figure 6B).

#### 3.7.2 Dark-Adapted

Combined rod-cone function was measured by the 3.0 SCR, and isolated rod function was measured using the 0.01 dim flash. There was no persistent significant difference between hypertonic buffer-injected, isotonic buffer-injected, or untreated eyes in this setting, suggesting minimal rod rescue following injections (Figure 6C-D).

These findings suggest that the injection of either isotonic or hypertonic buffer significantly improves cone-specific signaling pathways measured on ERG, and this beneficial effect persisted over the course of 5 months. In addition, these data suggest that the tonicity of the buffer introduced is a significant factor in cyst reduction and in ERG improvement, as more notable improvements were observed in eyes that received the hypertonic buffer compared to those that received the isotonic buffer.

### 3.8 Peak cyst severity on OCT is negatively associated with cone function on ERG at 5 MPI

Hypertonic buffer-injected eyes had both the lowest cyst severity and the greatest ERG improvements in cone dominant pathways. To explore this finding, we investigated whether cyst severity is truly associated with a long-term impairment in electrical signaling of the retina. Peak cyst severity, the highest cyst area reached by the eye, was graphed against ERG metrics at 5 MPI, and included hypertonic buffer, isotonic buffer, and untreated eyes. As part of the natural disease course in XLRS, retinal cysts hit a local maxima of severity between 2 and 4 months of age. The comparison of peak cyst severity to 5 MPI ERG amplitudes investigates whether the degree of cyst severity during critical times is correlated with long-term electrical function in the retina. By looking at long-term outcomes, we are able to see if the reduction of cysts could influence long-term retinal function beyond any short-term effect from bringing the retinal layers closer together. Additionally, at 5 MPI there are little to no cysts present in uninjected mice based on natural history and could not explain the variation in ERG amplitudes at that time.

In light-adapted, cone-dominant, ERG metrics we found ERG amplitudes are negatively correlated with peak cyst severity in both the 5 Hz flicker and 3.0 flash (*5 Hz flicker: p=0*.*0002;* Figure 7A) (*3*.*0 flash b-wave: p=0*.*016;* Figure 7B). These findings suggest greater cyst area has a more significant impact on impairing cone signaling than rod. Whether or not it is due to improved cone survival or improved cone functioning was unknown.

**Figure 7.**
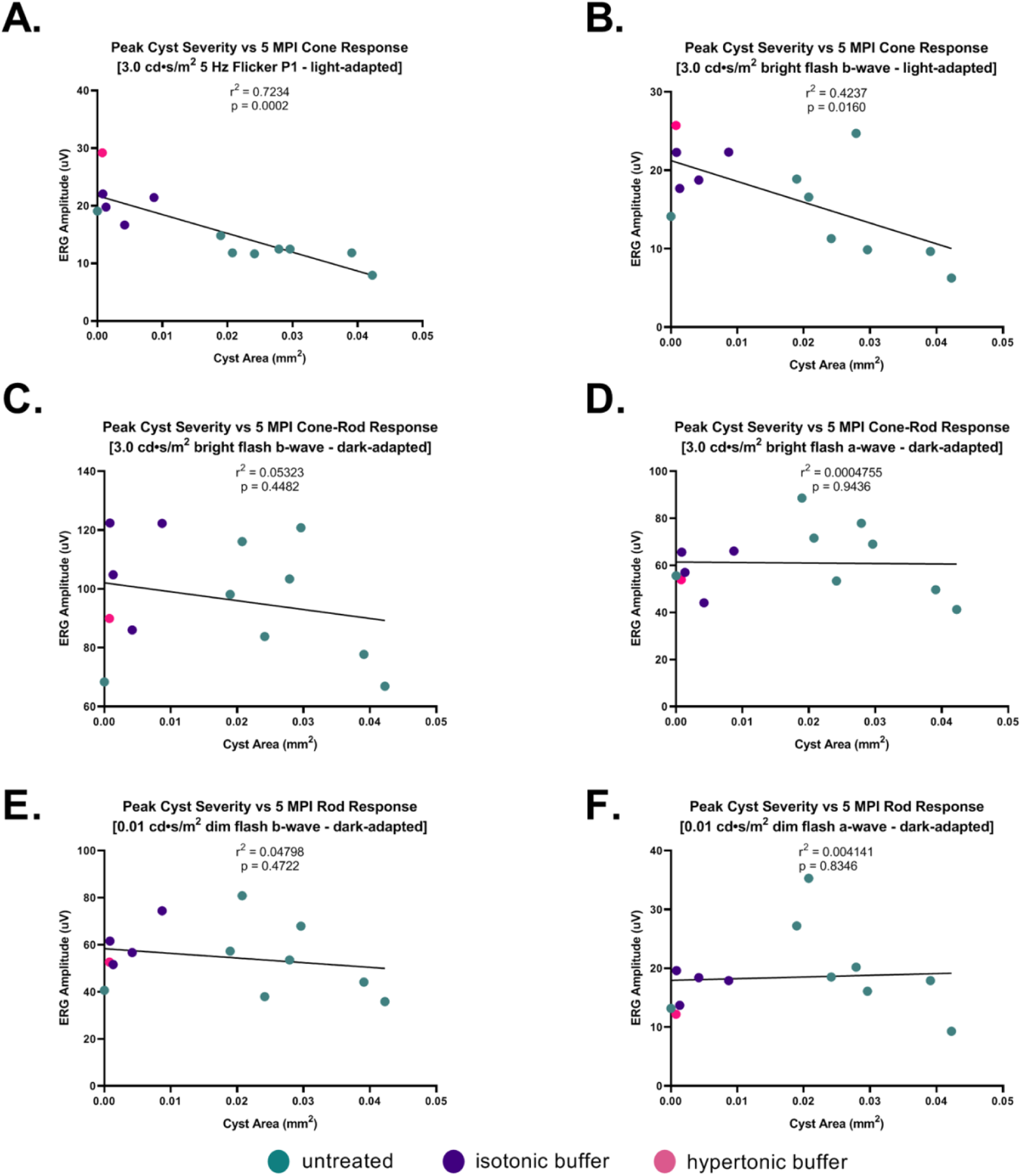
(A-F): Peak cyst severity on OCT is negatively associated with cone function on ERG at 5 MPI. Peak cyst severity of hypertonic buffer-injected, isotonic buffer-injected, and untreated eyes was determined by comparing the cyst area measurements of each eye at 1, 2, and 5 MPI. The cyst area of each eye representing their peak severity was correlated with its ERG amplitudes at 5 MPI. Each dot is representative of one eye. A negative correlation was found between cyst area and ERG amplitudes in the light-adapted 5 Hz Flicker (*p=0*.*0002*) and 3.0 cd•s/m^2^ bright flash (p= (A, B). Little to no correlation was found between peak cyst area and ERG amplitudes in dark-adapted 0.01 cd•s/m^2^ or 3.0 cd•s/m^2^ tests (C, D, E, F). These findings suggest increasing cyst area has greater impact on impairing cone electrical function, than rod photoreceptors.

On dark-adapted, rod-dominant ERG metrics, there is no correlation between peak cyst area in either the a-wave or b-wave metrics (*3*.*0 SCR b-wave: p=0*.*4482;* Figure 7C) (*3*.*0 SCR a-wave: p=0*.*9436;* Figure 7D) (*0*.*01 dim flash b-wave: p=0*.*4722;* Figure 7E) (*0*.*01 dim flash a-wave: p=0*.*8346;* Figure 7F). Cyst severity, or the reduction thereof, does not appear to influence long-term rod signaling.

To determine whether the improved cone amplitudes were associated with an increased number of cone photoreceptor cells, we used an immunofluorescence assay to quantify cone density in eyes injected with hypertonic buffer compared to untreated eyes.

### 3.9 Increased cone density at 6 months old suggests potential cone preservation mechanism following injection of hypertonic buffer

ONL thickness measurements, which are generally used to measure photoreceptor cell survival, showed no difference in thickness measurements between treated and untreated groups (Figure 5B). The overall stability in the ONL suggests rods survive long-term, regardless of cyst severity. In the setting of a low overall percentage of cones in the mouse retina (2.8% of total photoreceptors), the impact of the injections on cone survival may not be accurately determined by measuring outer nuclear layer thickness on conventional OCTs (22).

To reconcile improved ERG findings with the nonsignificant ONL findings, immunohistochemistry was performed to more precisely measure cone cell survival. Retinal cryosections were stained with biotinylated-PNA to quantify cone outer segments. These sections were then imaged, quantified, and averaged to give the output of cones per 100 μm of the retina. Hypertonic buffer-injected eyes had a mean of 20.95 cones per 100 μm, a significantly higher cone density than untreated *Rs1*-KO eyes with a mean of 13.42, suggesting improved cone survival compared to untreated eyes *(hypertonic buffer vs. Untreated: p=0*.*0147, Figure 8B)*.

**Figure 8.**
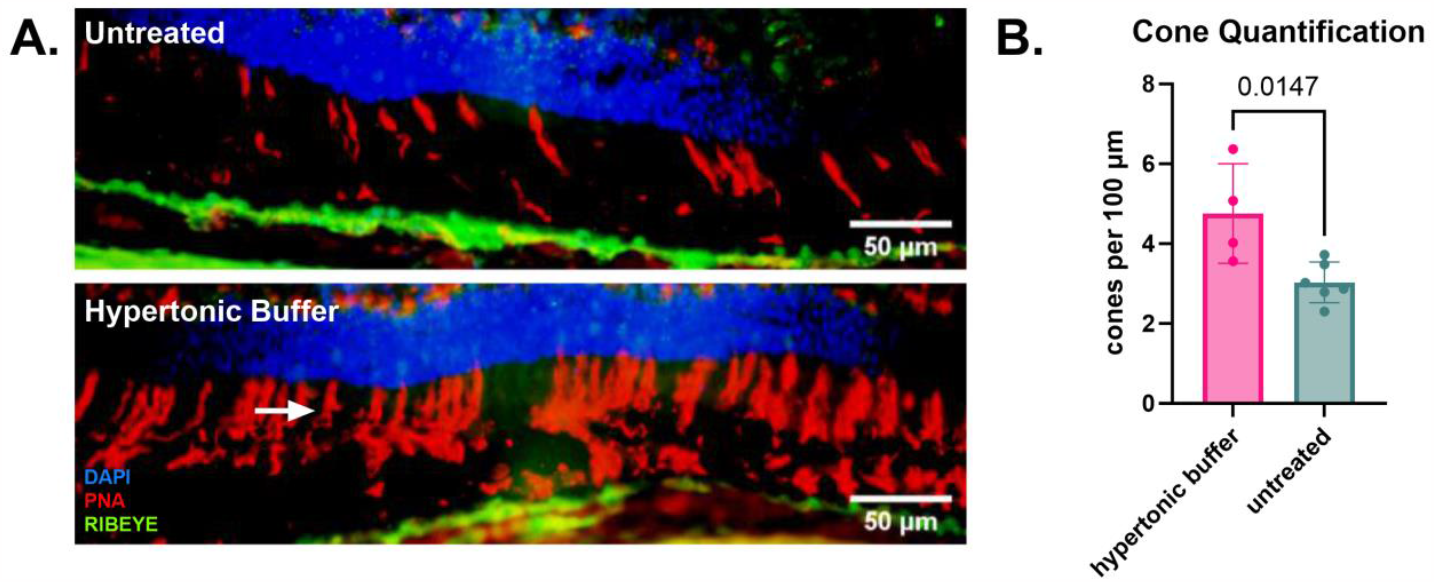
(A-B): Increased cone density at 6 months old suggests potential cone preservation mechanism following injection of hypertonic buffer. Representative immunofluorescent staining (RED) of cone outer segments, demonstrating increased density of cones in hypertonic buffer-injected eyes compared to untreated eyes with sparse cone staining (A). Retinal sections were collected from untreated *Rs1*-KO eyes and hypertonic buffer-injected eyes at approximately 6 months of age. Sections were stained by PNA to visualize cone outer segments. The number of cone outer segments per 100 μm of the retina was quantified by three masked participants, and averaged (cone outer segment indicated by white arrow). Green channel for RIBEYE staining of synapses. Each data point is representative of one eye. Hypertonic buffer-injected eyes show significantly higher cone density per 100 μm than untreated *Rs1*-KO eyes (*p=0*.*0147*) (B).

Together, this data shows that the subretinal injection of a buffer solution partially ameliorated intra-retinal cysts and lead to improved amplitudes in cone-specific ERGs as well as in functional day vision. The beneficial effect of the buffer solution after subretinal administration was partially mediated by the osmolarity of the buffer solution.

## 4 Discussion

XLRS gene therapy, both in mice and human trials, is currently underway, however, human trials have faced numerous challenges due to the development of adverse ocular immune reactions in some participants (9, 10). The primary goal of our study was to investigate whether low-titer gene therapy could circumvent immune reactions. We found that delivering a 2 × 10^9^ vg/mL dose of AAV2/4-EF1α-*RS1* subretinally in *Rs1*-KO mice did not cause meaningful rescue of functional metrics when compared to the metrics of a sham injection group, which received the hypertonic buffer injection alone. Because higher titers delivered using AAV2/4 under the CMV promoter subretinally and intravitreally have been shown to rescue the retinal phenotype (23), these results indicate that sufficient retinoschisin expression is required to achieve the highest efficacy.

Considered together with the data from the “sham” hypertonic buffer injections, it suggests that the efficacy of this very low dose vector may have been due solely to its diluent buffer.

Throughout our study, we observed that eyes that received subretinal injections of the hypertonic buffer consistently had less severe cysts and higher ERG amplitudes compared to the uninjected eyes of *Rs1*-KO mice. Those eyes that received hypertonic buffer injections performed comparably to our low dose gene therapy cohort which received the vector in the same buffer, even exhibiting a more robust effect at later time points when the gene therapy cohort had lost significance. Reduction of cysts for short periods following subretinal injection of a buffer has been reported in this model previously, but not explored thoroughly (19). In the prior report, retinal detachment was hypothesized to reduce cysts. The unexpected long term beneficial effects on XLRS phenotypes including ERG, VGSA, and photoreceptor survival after the subretinal injection of hypertonic buffer has not been previously reported.

Notably, the injection of hypertonic buffer alone attenuated the severity of cysts at 3 weeks, and this effect persisted until at least 2 months post injection. An overall reduction of cysts was apparent at 5 MPI across all groups, consistent with the waning of cysts that occurs during disease progression.

Interestingly, a concomitant improvement in cone-specific ERG amplitudes was observed in hypertonic buffer-treated eyes compared to those of uninjected eyes. Since the presence of cysts could interfere with bipolar cell function and impede electrical transduction, the improvements seen in ERGs could be a consequence of the attenuation of cyst severity in the retina.

Paradoxically, the improvements in cone-specific ERGs persisted despite age-related cyst resolution in both groups, and hypertonic buffer-injected eyes continued to possess higher ERG amplitudes at 5 MPI. Age-related cyst resolution in this model accompanies a blurring of the laminations on OCT, and may be associated with a new phase of retinal dysfunction. The thicknesses of the ONLs were not different between these groups of mice, so the reason behind the functional improvements seen in hypertonic buffer-injected eyes long after cysts have naturally resolved was unclear.

Since cones are reported to represent only 2.8 percent of photoreceptors in mouse retinas, it is likely that the effect of the injection on cone cell survival would not be reflected by ONL thickness measurements (22). We used immunohistochemistry to quantify the cone photoreceptor cells in eyes that received injections of buffer compared to in uninjected eyes. When we examined cone density using an immunofluorescent assay, we observed significantly higher cone density per 100 μm in hypertonic buffer-injected eyes than in untreated controls. An increase in overall cone density likely explains the improvement in ERG amplitudes seen in light-adapted settings and improved visual function in the light. This finding suggests a potential cone-preserving mechanism is associated with the subretinal introduction of hypertonic buffer. The relationship between cyst reduction and enhanced cone survival has not been previously reported and could have significant clinical implications.

Managing XLRS symptoms using topical agents such as brinzolamide has been found to reduce cysts along with a beneficial increase in visual function (6-8). However, whether managing cysts in XLRS could have long-term benefits is not clear. Our study provides compelling evidence that the attenuation of cysts may promote the survival of cones in the retina, and lead to better electrical cone signaling and functional vision in the long-term.

The effects of a hypertonic buffer subretinal-injection on improving the function and structure of the retina have not been previously reported. Previous studies propose that retinoschisin 1 (*RS1*) might have additional functions outside of its structural function and play a role in governing fluid distribution across retinal layers, potentially by engaging with Na/K-ATPase channels via a binding mechanism (20, 21). In the absence of a functional protein, this pathway has the potential to become dysregulated, resulting in the formation of fluid filled retinal cysts and creating a spatial separation of bipolar and photoreceptor cells. Despite the incomplete understanding of the mechanism, this phenomenon could provide an explanation for the observed benefits of carbonic anhydrase inhibitors (CAIs) in mitigating cyst development and enhancing visual function among XLRS patients (2).

These data provided the rationale behind our investigation whether tonicity of the buffer could affect fluid movement out of cysts and resolve schisis.

ERG and OCT experiments were repeated with an isotonic buffer to determine if this effect was influenced by the tonicity of the solution versus the mechanical action of the subretinal injection. We found that the isotonic buffer-injected cohorts did not perform as well as the hypertonic buffer-injected eyes. Though isotonic buffer-treated eyes had reduced cysts, similar but slightly greater than hypertonic buffer-injected eyes, they experienced lower b-wave amplitudes in light-adapted ERGs.

These experiments indicate that the tonicity of the injected solution plays a role in the reduction of cysts and function of the retina after injection.

Currently, our findings suggest that the injection of a buffer that is hypertonic to the physiological environment alleviates intraretinal cysts, promotes the survival of cone photoreceptors, and preserves retinal function in a mouse model of XLRS. Furthermore, these data suggested that managing the severity of cysts can have a long-term impact on the health of the retina. To examine this notion, we analyzed the relationship between peak cyst severity of each animal with their long-term functional outcome. When we measured peak cyst severity against endpoint ERG amplitudes, we observed a negative correlation between the peak severity of cysts and ERG amplitudes in light-adapted ERG settings. This correlation was seen regardless of the treatment cohort (untreated, hypertonic buffer-, and isotonic buffer-injected), suggesting that the light-adapted metrics, and thus cone function and or survival, are closely related to severity of cysts. In the setting of greater cyst severity, cone function is more attenuated than rod function. This also suggests that long term cone function can be predicted by the degree of cyst reduction achieved by treatment. These results suggest the successful management of cysts can impact the long-term health and function of the retina.

This study has provided evidence that the management of cysts through an osmolarity-related mechanism can have beneficial effects. Improvements in cone function and survival, cyst severity, and light-adapted visual function after hypertonic buffer-injection suggest tonicity is a mediator of this effect. We speculate that this effect is due to the shifting of fluid out of the retinal space, where cysts form, and into the subretinal space where the choroidal vasculature can reabsorb the cystic fluid.

Aquaporins (AQP) represent a family of water-transporting channels with a pivotal role in upholding water homeostasis across various organ systems. They have been previously identified in the inner nuclear layer of the retina, where studies suggest they have an involvement in regulating retinal water balance (24). When AQP was knocked out in mouse models, electroretinograms (ERGs) demonstrated decreased b-wave amplitudes and delayed latency in a series of ERGs with increasing light intensity (24). These findings are parallel to our observations in the *Rs1*-KO mouse natural history and underscore the connection between the electrical function of the eye and the fluid homeostasis of the retina. Systemic infusions of hyperosmolar fluid have been shown to reduce edema in other organs, like the brain, in the setting of acute cerebral injury (25). With this precedent, strategies leveraging fluid exchange to promote the resolution of cysts in edematous retinal conditions merit further exploration.

Although the exact mechanism is unknown, we have shown that injection of a buffer into the subretinal space leads to reduced cyst severity in the *Rs1*-KO mouse and that lower peak cyst severity is associated with improved electrical function, improvements in light-adapted visual acuity, and greater preservation of cones. This effect is partially modulated by the tonicity of the buffer and is most prominent in eyes injected with a hypertonic buffer. These findings have the potential to open new doors in the development of therapies for patients with XLRS and could shine a light on how fluid balance influences cyst resolution and disease severity.

## 7 Acknowledgments and Funding

We would like to thank Dalyz Ochoa and Budd Tucker Ph.D. for providing the AAV2/4-EF1α-*RS1* vector for this study. We would also like to thank Paul Sieving M.D., Ph.D. for providing us with the *Rs1*-KO mouse line.

This work was generously supported by the Chakraborty Family Foundation and the Keech Professorship.

## References

1. Strupaite R, Ambrozaityte L, Cimbalistiene L, Asoklis R, Utkus A. X-linked juvenile retinoschisis: phenotypic and genetic characterization. Int J Ophthalmol. 2018;11(11):1875–8.

2. Molday RS, Kellner U, Weber BH. X-linked juvenile retinoschisis: clinical diagnosis, genetic analysis, and molecular mechanisms. Prog Retin Eye Res. 2012;31(3):195–212.

3. Sergeev YV, Vitale S, Sieving PA, Vincent A, Robson AG, Moore AT, et al. Molecular modeling indicates distinct classes of missense variants with mild and severe XLRS phenotypes. Hum Mol Genet. 2013;22(23):4756–67.

4. Sieving PA, MacDonald IM, Hoang S. X-Linked Congenital Retinoschisis. In: Adam MP, Mirzaa GM, Pagon RA, Wallace SE, Bean LJH, Gripp KW, et al., editors. GeneReviews((R)). Seattle (WA)1993.

5. Wolfensberger TJ, Dmitriev AV, Govardovskii VI. Inhibition of membrane-bound carbonic anhydrase decreases subretinal pH and volume. Doc Ophthalmol. 1999;97(3-4):261–71.

6. Apushkin MA, Fishman GA. Use of dorzolamide for patients with X-linked retinoschisis. Retina. 2006;26(7):741–5.

7. Scruggs BA, Chen CV, Pfeifer W, Wiley JS, Wang K, Drack AV. Efficacy of topical brinzolamide in children with retinal dystrophies. Ophthalmic Genet. 2019;40(4):350–8.

8. Thobani A, Fishman GA. The use of carbonic anhydrase inhibitors in the retreatment of cystic macular lesions in retinitis pigmentosa and X-linked retinoschisis. Retina. 2011;31(2):312–5.

9. Cukras C, Wiley HE, Jeffrey BG, Sen HN, Turriff A, Zeng Y, et al. Retinal AAV8-RS1 Gene Therapy for X-Linked Retinoschisis: Initial Findings from a Phase I/IIa Trial by Intravitreal Delivery. Mol Ther. 2018;26(9):2282–94.

10. Pennesi ME, Yang P, Birch DG, Weng CY, Moore AT, Iannaccone A, et al. Intravitreal Delivery of rAAV2tYF-CB-hRS1 Vector for Gene Augmentation Therapy in Patients with X-Linked Retinoschisis: 1-Year Clinical Results. Ophthalmol Retina. 2022;6(12):1130–44.

11. Reichel FF, Peters T, Wilhelm B, Biel M, Ueffing M, Wissinger B, et al. Humoral Immune Response After Intravitreal But Not After Subretinal AAV8 in Primates and Patients. Invest Ophthalmol Vis Sci. 2018;59(5):1910–5.

12. Kjellstrom S, Bush RA, Zeng Y, Takada Y, Sieving PA. Retinoschisin gene therapy and natural history in the Rs1h-KO mouse: long-term rescue from retinal degeneration. Invest Ophthalmol Vis Sci. 2007;48(8):3837–45.

13. Wiley LA, Burnight ER, Kaalberg EE, Jiao C, Riker MJ, Halder JA, et al. Assessment of Adeno-Associated Virus Serotype Tropism in Human Retinal Explants. Hum Gene Ther. 2018;29(4):424–36.

14. Zeng Y, Takada Y, Kjellstrom S, Hiriyanna K, Tanikawa A, Wawrousek E, et al. RS-1 Gene Delivery to an Adult Rs1h Knockout Mouse Model Restores ERG b-Wave with Reversal of the Electronegative Waveform of X-Linked Retinoschisis. Invest Ophthalmol Vis Sci. 2004;45(9):3279–85.

15. McCulloch DL, Marmor MF, Brigell MG, Hamilton R, Holder GE, Tzekov R, et al. Erratum to: ISCEV Standard for full-field clinical electroretinography (2015 update). Doc Ophthalmol. 2015;131(1):81–3.

16. Hsu Y, Bhattarai S, Thompson JM, Mahoney A, Thomas J, Mayer SK, et al. Subretinal gene therapy delays vision loss in a Bardet-Biedl Syndrome type 10 mouse model. Mol Ther Nucleic Acids. 2023;31:164–81.

17. Bush RA, Zeng Y, Colosi P, Kjellstrom S, Hiriyanna S, Vijayasarathy C, et al. Preclinical Dose-Escalation Study of Intravitreal AAV-RS1 Gene Therapy in a Mouse Model of X-linked Retinoschisis: Dose-Dependent Expression and Improved Retinal Structure and Function. Hum Gene Ther. 2016;27(5):376–89.

18. Salma Hassan YH, Sara K. Mayer, Jacintha Thomas, Aishwarya Kothapalli, Megan Helms, Sheila A. Baker, Joseph G. Laird, Sajag Bhattarai, Arlene V. Drack. A visually guided swim assay for mouse models of human retinal disease recapitulates the multi-luminance mobility test in humans. [in press]. In press 2023.

19. Luna G, Kjellstrom S, Verardo MR, Lewis GP, Byun J, Sieving PA, et al. The effects of transient retinal detachment on cavity size and glial and neural remodeling in a mouse model of X-linked retinoschisis. Invest Ophthalmol Vis Sci. 2009;50(8):3977–84.

20. Molday LL, Wu WWH, Molday RS. Retinoschisin (RS1), the Protein Encoded by the X-linked Retinoschisis Gene, Is Anchored to the Surface of Retinal Photoreceptor and Bipolar Cells through Its Interactions with a Na/K ATPase-SARM1 Complex*. Journal of Biological Chemistry. 2007;282(45):32792–801.

21. Plössl K, Royer M, Bernklau S, Tavraz NN, Friedrich T, Wild J, et al. Retinoschisin is linked to retinal Na/K-ATPase signaling and localization. Mol Biol Cell. 2017;28(16):2178–89.

22. Jeon CJ, Strettoi E, Masland RH. The major cell populations of the mouse retina. J Neurosci. 1998;18(21):8936–46.

23. Scruggs BA, Bhattarai S, Helms M, Cherascu I, Salesevic A, Stalter E, et al. AAV2/4-RS1 gene therapy in the retinoschisin knockout mouse model of X-linked retinoschisis. PLoS One. 2022;17(12):e0276298.

24. Li J, Patil RV, Verkman AS. Mildly abnormal retinal function in transgenic mice without Müller cell aquaporin-4 water channels. Invest Ophthalmol Vis Sci. 2002;43(2):573–9.

25. Zhang H, Liu J, Liu Y, Su C, Fan G, Lu W, et al. Hypertonic saline improves brain edema resulting from traumatic brain injury by suppressing the NF-kappaB/IL-1beta signaling pathway and AQP4. Exp Ther Med. 2020;20(5):71.

